# Rubisco from two ecotypes of *Plantago lanceolata* L. that are native to sites differing in atmospheric CO_2_ levels

**DOI:** 10.1101/2024.07.13.603406

**Authors:** Xiaoxiao Shi, Nathan M. Hannon, Arnold J. Bloom

## Abstract

Rubisco, the most prevalent protein on the planet, initiates the conversion of CO_2_ into carbohydrates during photosynthesis. Responses of this process to atmospheric CO_2_ fluctuations daily, seasonally, and over millennia is still poorly understood. We have hypothesized that higher plants maintain carbon-to-nitrogen homeostasis by adjusting the balance of magnesium and manganese in chloroplasts to alter their relative carbon fixation and nitrogen assimilation rates. The following study examined the influence of magnesium and manganese on rubisco carboxylation and oxygenation in protein purified from two ecotypes of *Plantago lanceolata*: one adapted to the high atmospheric CO_2_ that occurs near a natural CO_2_ spring and the other adapted to more typical CO_2_ atmospheres that occur nearby. The plastid DNA coding for the large unit of rubisco were similar in both ecotypes. The kinetics of rubiscos from the two ecotypes differed more when they were associated with manganese than magnesium. Specificity for CO_2_ over O_2_ (*S*_c/o_) for rubiscos from both ecotypes were higher when the enzymes were bound to magnesium than manganese. This disparity may account for the adaptation of this species to different CO_2_ environments.

## Introduction

Global atmospheric CO_2_ will increase from its current concentration of 421 ppm to somewhere between 450 and 1130 ppm by the end of the century (IPCC, 2021). Experiments on the responses of higher plants to rising CO_2_ subject them to atmospheres containing higher than 550 ppm CO_2_ (Broberg *et al*., 2019; Tcherkez *et al*., 2020), a concentration that is 40% above the current ambient. Such treatments accelerate carbon fixation, inhibit photorespiration (Cousins and Bloom, 2004), decrease protein concentrations by over 6% (Medek *et al*., 2017; Myers *et al*., 2014; Taub *et al*., 2008), and increase plant carbon-to-nitrogen ratio (C/N) by 20% (Butterly *et al*., 2015; Krämer *et al*., 2022; Sardans *et al*., 2012; Wang *et al*., 2019; Ziska *et al*., 2016).

Central to these responses is rubisco (ribulose-1,5-bisphosphate carboxylase-oxygenase), the enzyme that constitutes 50% of the protein in a leaf, 20% of the protein on Earth, and 3% of the global biomass (Bar-On and Milo, 2019; Ellis, 1979). Rubisco catalyzes several competing reactions (Bloom and Lancaster, 2018): One reaction is the carboxylation of ribulose 1,5-bisphosphate (RuBP) to two molecules of 3-phosphoglyceric acid (3PGA) that initiates C_3_ carbon fixation, the foundation of primary productivity. A second reaction is the oxygenation of RuBP to one molecule of 3PGA and one molecule of 2PG that initiates photorespiration, a process generally considered to dissipate more than 25% of the energy in C_3_ plants as waste heat (Smith *et al*., 2024; South *et al*., 2019). Photorespiration, however, enhances assimilation of nitrate (NO_3_^−^) and sulfate (SO_4_^−^) into amino acids in plant shoots (Bloom, 2015a, b, 2019; Bloom *et al*., 2010; Bloom *et al*., 2014; Bloom and Kameritsch, 2017; Bloom *et al*., 2020; Bloom and Lancaster, 2018; Bloom and Plant, 2021; Bloom *et al*., 2012; Bloom *et al*., 2002; Carlisle *et al*., 2012; Dietterich *et al*., 2015; Easlon and Bloom, 2013; Foyer *et al*., 2009; Myers *et al*., 2014; Rachmilevitch *et al*., 2004; Rubio-Asensio and Bloom, 2017; Rubio-Asensio *et al*., 2015; Smart and Bloom, 2001; Smart *et al*., 1998). Therefore, plant protein concentrations, growth, and yields decrease when elevated CO_2_ atmospheres inhibit photorespiration for long periods (Bloom, 2015a; Bloom *et al*., 2014; Bloom and Plant, 2021; Myers *et al*., 2014).

A handful of studies conducted more than four decades ago indicated that binding of rubisco to Mg^2+^ instead of Mn^2+^ accelerates RuBP carboxylation and inhibits RuBP oxygenation, but the data were highly variable (Christeller, 1981; Christeller and Laing, 1979; Jordan and Ogren, 1983; Martin and Tabita, 1981; Wildner and Henkel, 1979). These studies purified rubisco via ammonium sulfate precipitation followed by centrifugation, a process that can adversely influence enzyme structure and activation (Iñiguez *et al*., 2021; Wingfield, 1998). In a recent study (Shi *et al*., 2024), we used fast protein liquid chromatography to purify rubisco from five model C_3_ species while preserving the metal-binding characteristics of the native protein (Barnett *et al*., 2012; Hagège *et al*., 2015; Montes-Bayón *et al*., 2016) and developed new methods to assess rubisco carboxylation and oxygenation that are indifferent to the presence of Mg^2+^ or Mn^2+^ (Shi *et al*., 2024). The maximum velocity of carboxylation (*V*_cmax_) was faster and the Michaelis constants of rubisco for CO_2_ (*K*_c_) was greater when rubisco was bound to Mg^2+^ rather than Mn^2+^. Both *S*_c/o_ (rubisco specificity for CO_2_ over O_2_) and *V*_cmax_/*V*_omax_ were greater when the enzyme was bound to Mg^2^ rather than Mn^2+^ (Shi *et al*., 2024).

*Plantago lanceolata* L. may not be a model plant species, but its leaves have been widely used in herbal medicines (Drava et al. 2019, Gonçalves and Romano 2016, Wichtl 2004) and food preparations (Guarrera and Savo 2016). Research has shown that the root fractions of *P. lanceolata* L. possess antibacterial properties as demonstrated through Phytochemical Analysis by Gas Chromatography-Mass Spectrometry *in vitro* (Rahamouz-Haghighi et al. 2022). Furthermore, *P. lanceolata* leaf extract has been found to selectively inhibit the proliferation of CAL51 triple-negative breast cancer cell proliferation (Alsaraf et al. 2019) and have anti-obesity effects on mice *in vivo* (Yoshida et al., 2013). Several flavonoids present in the inflorescences and leaves of *P. lanceolata* have been discovered to have anti-allergic and anti-inflammatory effects (Budzianowska et al. 2022, Murai et al., 1995, Antunes-Ricardo et al. 2015).

The following study compared the kinetics of rubisco purified from two ecotypes of *Plantago lanceolata* L.: one collected near a CO_2_ spring has experienced an average daytime concentration of 791 ppm CO_2_ for many generations, and the other collected 200 m from the spring has experienced ambient CO_2_ concentrations of about 421 ppm (Saban *et al*., 2019; Saban *et al*., 2020; Watson-Lazowski *et al*., 2016). We also extracted plastid DNA from the two *P. lanceolata* ecotypes and compared the sequences for the large subunit of rubisco (*rbcL*) to determine if multi-generational exposures to different atmospheric CO_2_ levels might result in genetic adaptations in rubisco.

## Materials and methods

### Plant growth conditions

Prof. G. Taylor provided us with seeds from of *Plantago lanceolata* L. collected in May 2008 from nine plants growing in naturally elevated CO_2_ atmospheres near a CO_2_ spring at Bossoleto, Italy (Lat. 43°17’, Long. 11°35’) and at a nearby (ca. 200 m apart) ambient CO_2_ control site (Saban *et al*., 2020). Seeds obtained from the two sites were grown for one generation in the glasshouse at the University of Southampton and then crossed within maternal families to standardize parental effects. We planted the two ecotypes into 5 × 5 cm containers containing Sunshine Mix 4 (Sungro, Agawam, MA). The plants grew in a controlled environmental chamber for 21 d under 450 ppm or 750 ppm CO_2_, 60% relative humidity (light/dark), air temperature 25°/18°C (light/dark), and 16 h of 500 μmol m^−2^ s^−1^ PPFD at canopy height.

### Rubisco extraction and purification

We collected about 40 g of leaves and froze them in liquid N_2_. Manual grinding with a mortar and pestle lysed the plant cells to a fine powder. We extracted the fine powder in 100 ml of a buffer (50 mM Tris-HCl pH 7.4, 20 mM MgCl_2_, 20 mM NaHCO_3_, 0.1 mM Na_2_EDTA, 10% glycerol, 50 mM mercaptoethanol, and 1 mM PMSF), filtered the extract through four layers of Miracloth, centrifuged it at 12100 × *g* at 4°C for 15 min to clarify it, and passed the supernatant through a 0.22 μm syringe filter before loading it onto an FPLC column.

We purified rubisco using an ENrich™ SEC (Size Exclusion Column) 650 10 × 300 Column (BioRad, Hercules, CA) followed by a HiScale 16/20 6 mL SOURCE 30Q column (GE Healthcare Life Sciences, Pittsburgh, PA) on a NGC™ FPLC system (BioRad, Hercules, CA). All buffers included 2 mM dithiothreitol to prevent intermolecular disulfide bond formation. Pre-equilibration of the size exclusion column involved eluting five column volumes (CV) of 85% buffer A (50 mM Tris-HCl, 1 mM EDTA, 0.1 mM PMSF, pH 7.4) and 15% buffer B (50 mM Tris-HCl, 1 M NaCl 1 mM EDTA, 0.1 mM PMSF, pH 7.4). We loaded the protein sample onto the column and eluted it with 2 CV of 15% buffer B. UV absorption at 280 nm confirmed the protein peak. We (*a*) concentrated the protein solution to around 1 mM, (*b*) exchanged the buffer into buffer A, (*c*) filtered the sample, (*d*) loaded it onto a SOURCE 30Q column pre-equilibrated with 5 CV of buffer A, (*e*) eluted the protein with 50% buffer B with a linear gradient over 10 CV, (*f*) pooled the protein peak at 280 nm, (*g*) exchanged the buffer into buffer A with the addition of 20% glycerol, (*h*) and stored the protein solution at -80°C. We checked the purity of the protein on SDS-PAGE gels, Western Blots, and an Evolution 201 UV/Vis spectrometer coupled with an Evolution 1-cell Peltier temperature control system (Thermo Fisher Scientific, Waltham, MA).

### Rubisco carboxylation colorimetric reaction

We developed a colorimetric assay for rubisco carboxylation that is accurate in the presence of Mg^2+^ or Mn^2+^ (Shi *et al*., 2024):

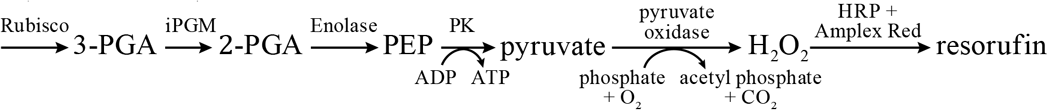

To estimate *v*_c_ and *V*_cmax_, we assessed the production of 3PGA in the presence of high levels of CO_2_ by iPGM (cofactor-Independent PhosphoGlycerate Mutase) conversion of 3PGA to 2PGA (Raverdy *et al*., 2007; Zhang *et al*., 2004), followed by a commercial assay kit (2-phosphoglycerate Assay kit, Abcam ab174097) that converts 2PGA into PEP, then pyruvate, H_2_O_2_, and resorufin. Absorption at 570 nm monitored resorufin production. This assay can detect 2PGA levels below 20 pmol and works in the presence of either metal cofactor.

In more detail, we activated rubisco before the assays as follows. We added an assay buffer containing 20 mM Tris-HCl, 1 mM EDTA, 100 mM Triethanolamine at pH 7.8 to one 5 mL tube; added 1 or 2 μL of rubisco (0.5 – 1 μM), 10 μL of 250 mM NaHCO_3_, 0.5 μL carbonic anhydrase (∼2.5 units), 5 μL of iPGM (10 μM), 4 μL of MgCl_2_ (final concentration 20 mM) or MnCl_2_ (5 mM); mixed thoroughly; and allowed the mixture to sit for 5 minutes. The final assay volume in the tube was 1750 μL. We started the reaction by adding 45 μL of 10 mM RuBP to the tube, split the mixture into five tubes, and stopped the reaction after 1, 2, 3, 4, or 5 mins by adding 0.5 N HCl to each tube. We added KOH to each reaction tube to adjust the pH to about pH 7.8 and added to the tubes equal amounts of freshly mixed 2PGA colorimetric cocktail that we prepared from a 2-phosphoglycerate assay kit (ab174097, Abcam, Cambridge, MA), mixed well, and moved the tubes to an opaque box. We measured OD_570nm_ after 40 min for each of the tubes and calculated the 2PGA concentration in each reaction tube based on a calibration curve determined by adding specified quantities of a 1 mM 2PGA standard solution that the assay kit provided.

The carboxylation turnover rate *v*_c_ = ½ (resorufin production rate – *v*_o_), where O_2_ optode measurements of O_2_ depletion estimated *v*_o_ as detailed below.

We estimated *V*_cmax_ from the Michaelis-Menten equation:

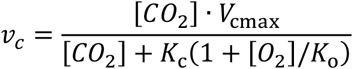

and the relative specificity *S*_c/o_ from:

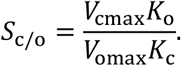

We conducted separate calibration runs in the presence of the buffer and Mg^2+^ or Mn^2+^ in the same concentrations used in the rubisco reactions (**Fig. S1**) and calculated the ratio between the slopes of the runs using only the 2PGA buffer and the slopes of the runs containing rubisco. Linear regressions for each combination of plant and metal ion, which included a constant term for each run, expressed the change in 2PGA as a function of time.

### IPGM expression and purification

We transformed a 6xHis-tagged *C. elegans* nematode iPGM plasmid into BL21(DE3) competent cells and incubated them at 37°C overnight. We selected one colony for culturing at 37°C overnight in 50 mL LB media (peptone 10 g, yeast extract 5 g, NaCl 10 g L^−1^) and 50 μL 100 g mL^−1^ kanamycin stock solution. We added 10 mL of the small culture to 1 L of the same media and grew it until OD_600_ = 0.7. We added isopropyl β-D-1-thiogalactopyranoside (IPTG) to the culture flask to bring it to a final concentration of 0.2 mM and reduced the temperature and stirring speed to 30°C and 100 rpm, respectively. We grew the culture for 24 h, centrifuged it for 20 min at 4700 × *g*, and suspended the pellet in 30 mL buffer A (20 mM Tris-HCl, 2 mM DTT, pH 6.8). We stored the cell suspension at −80°C before purification.

We thawed 30 mL of the cell suspension, lysed the cells by adding 30 mg of lysozyme and 30 μg of DNAse, and passed the cells twice through a French press. We centrifuged the solution at 12100 × *g* for 60 min, concentrated the supernatant using Amicon Ultra Centrifuge Filters (EMD Millipore, Burlington, MA), and filtered the resulting solution through a 0.22 μm filter before loading it onto a column.

We pre-equilibrated a Bio-Scale Mini Profinity IMAC cartridge (Bio-Rad, Hercules, CA) with five column volumes (CV) of buffer A (50 mM Tris-HCl, 25 mM imidazole, 1 mM EDTA, 0.1 mM PMSF, pH 7.4), loaded the protein sample onto the column, washed the column with 5 CV of buffer A, and eluted the protein with a linear gradient over 10 CV of buffer B (50 mM Tris-HCl, 500 mM imidazole, 1 mM EDTA, 0.1 mM PMSF, pH 7.4), confirmed the protein peak by UV absorption at 280 nm, and loaded the fractions onto an SDS-PAGE gel. We stored the fractions with iPGM protein at –80°C.

### Rubisco oxygenation

A needle-type micro-optode OXF50-OI (PyroScience GmbH, Breman, Germany) on a FireSting O_2_ optical oxygen and temperature meter (FSO2-4) monitored changes in O_2_ concentration. We conducted oxygenation experiments under three sets of conditions at 25°C: (*a*) ambient (79% N_2_, 20.96% O_2_, and 0.04% CO_2_), (*b*) elevated CO_2_ (78.96% N_2_, 20.96% O_2_, and 0.08% CO_2_), and (*c*) reduced O_2_ (89% N_2_, 10.96% O_2_, and 0.04% CO_2_). Precision mass flow controllers (Apex Vacuum, Canton, GA) calibrated against soap bubble flowmeters mixed pure N_2_, O_2_, and CO_2_ to the desired concentrations. A non-dispersive Infrared Gas Analyser (Li-cor, Lincoln, NE) checked the CO_2_ concentration. We conducted a two-point calibration of the oxygen optode in both air and air-saturated water. The final assay volume in each tube was 3000 μL.

Before the oxygenation assay, we activated rubisco as follows. We added assay buffer (20 mM Tris-HCl, 1 mM EDTA, 10 mM NaHCO_3_, 100 mM triethanolamine, pH 7.8) to a covered, temperature-controlled, 4 mL quartz cuvette; added 10 μL of rubisco (0.5 – 1 μM), 50 μL of iPGM (10 μM), 5 μL carbonic anhydrase (∼25 units), and 40 μL of MgCl_2_ (final concentration 20 mM) or MnCl_2_ (5 mM); and mixed the contents of the cuvette thoroughly. We allowed the mixture to sit for 5 minutes before inserting the oxygen sensor and sparging for 1 minute with the gas mixture corresponding to the experiment’s conditions. After about 30 s, once the oxygen sensor reading stabilized, we started the reaction by adding 100 μL of 10 mM RuBP (pre-equilibrated with the same gas mixture) and began collecting data. We did not observe any apparent protein deactivation and degradation during more than 300 s; therefore, we estimated the oxygenation turnover rate *v*_o_ from the linear trend in oxygen consumption for at least 200 s divided by rubisco content.

Michaelis-Menten kinetics predicts the oxygenation turnover rate *v*_o_ to be:

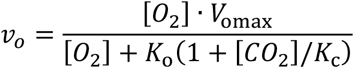

where *V*_omax_, *K*_o_, and *K*_c_ depend on species and the associated metal, Mn^2+^ or Mg^2+^, but are independent of [O_2_] and [CO_2_]. Values of *v*_o_ under multiple sets of conditions for [O_2_] and [CO_2_], therefore, provided estimates of the three parameters *V*_omax_, *K*_o_, and *K*_c_. For the reduced O_2_ experiments, we performed blank runs with RuBP and all other components but without rubisco. We calculated O_2_ consumption rates before and after adding RuBP for both the blank and actual runs and determined the O_2_ consumption rate as

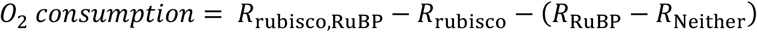

where *R* denotes the rate of change in oxygen levels in the presence or absence of rubisco and RuBP.

### Statistics

In general, we used R for Windows version v.4.3.3 and will provide the R code upon request. From the measured rates of rubisco reactions, kinetic parameters were calculated using a Bayesian method. Briefly, random sets of kinetic parameters were generated from a lognormal prior distribution based on the approximate range in which kinetic parameters for C_3_ plants have been calculated, and the relative likelihood of observing the values in our experiments was computed for each set of kinetic parameters.

The result of each experiment was assumed to be normally distributed with a fixed variance, combining the regression standard error in the measurements with the variance between the results of similar experiments. These likelihoods were used as weights to approximate a posterior distribution on the kinetic parameters from which we calculated posterior means and standard deviations of each parameter.

To determine the overall effects of Mg^2+^ and Mn^2+^ across the two ecotypes, we aggregated each kinetic parameter, adjusted by their respective standard errors, and used it as input to a two-way ANOVA, with metal ion and ecotype as variables.

### *rbcL* sequences

We isolated chloroplasts (Bloom and Kameritsch, 2017) from the two *P. lanceolata* ecotypes and extracted the plastid DNAs using the modified high salt method (Shi et al. 2012). A_260_/A_280_ ratios were verified by UV-Vis spectrophotometry, and samples with a ratio over 2 were sent for chloroplast genomic DNA sequencing (CD Genomics, Shirley, NY). After conducting the initial sample quality test, high-quality DNA samples were used to construct the library. The purified DNA samples were fragmented into 200-500 bp using Covaris S/E210 or Bioruptor. The overhangs resulting from the fragmentation were converted into blunt ends using T4 DNA polymerase, Klenow Fragment, and T4 Polynucleotide Kinase.

Subsequently, an ‘A’ base was added to the 3’ end of the blunt phosphorylated DNA fragments, followed by the ligation of adapters to the ends of the DNA fragments. The desired fragments were then purified through gel-electrophoresis, selectively enriched, and amplified by PCR. Index tags were introduced into the adapter during the PCR stage, and a library quality test was performed. Finally, the qualified library was used for sequencing using Illumina PE150 Sequencing Platform, and the generated data were utilized for downstream bioinformatics analysis. GetOrganelle (v1.7.7.1) was used to construct the final cpDNA sequences. Custom R code (v4.3.3) was used to search for fragments containing variants of the cpDNA sequences and determine their frequencies. For each identified variant, a sequence of five codons centered on the changed nucleotide(s) was chosen, and the number of fragments containing the original and variant sequences was compared to calculate its frequency in both ecotypes.

## Results

### Rubisco Kinetics

Rubisco purified from the ambient ecotype conducted both RuBP carboxylation and oxygenation at higher rates under all conditions than rubisco purified from the CO_2_ spring ecotype (**Figs. 1 & 2**). Rubisco from the ambient ecotype conducted significantly more carboxylation under 0.10% CO_2_ than 0.04% CO_2_, whereas rubisco from the spring ecotype was much less affected by CO_2_ enrichment (**Figs. 1 & 2**). Rubiscos from both ecotypes conducted significantly more RuBP carboxylation and oxygenation when bound to Mn^2+^ compared to Mg^2+^. (**Fig. 2**).

**Fig. 1.**
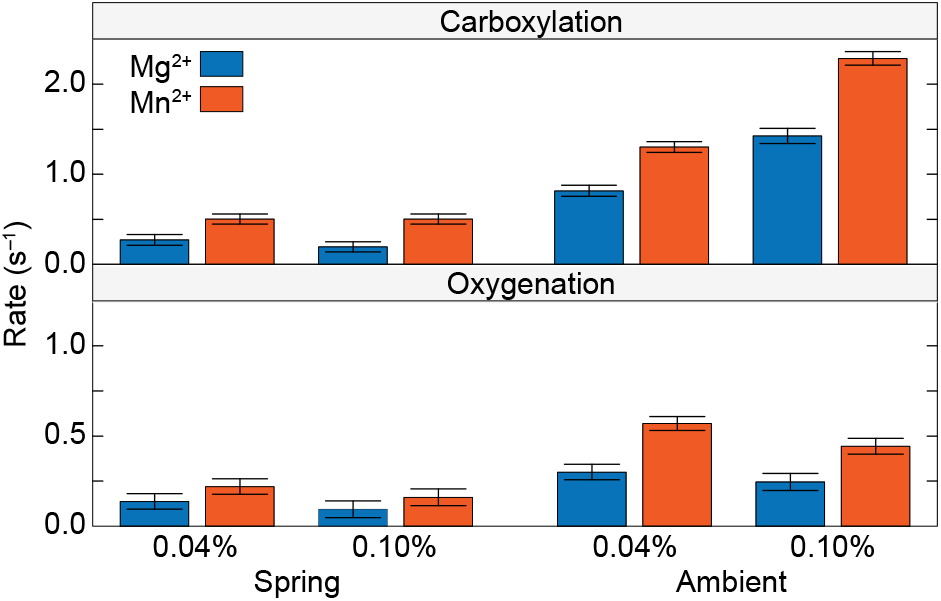
Rates of RuBP carboxylation and oxygenation in turnovers per second by rubisco purified from two ecotypes of *Plantago lanceolata* L. One ecotype (Spring) collected near a CO_2_ spring had experienced high CO_2_ for many generations, and the other ecotype collected nearby (Ambient) had experienced ambient CO_2_ atmospheres. The enzyme was bound to Mg^2+^ or Mn^2+^ and exposed to 0.04% or 0.10% CO_2_ atmospheres. Shown are the mean ± SE, n= 4 – 10.

**Fig. 2.**
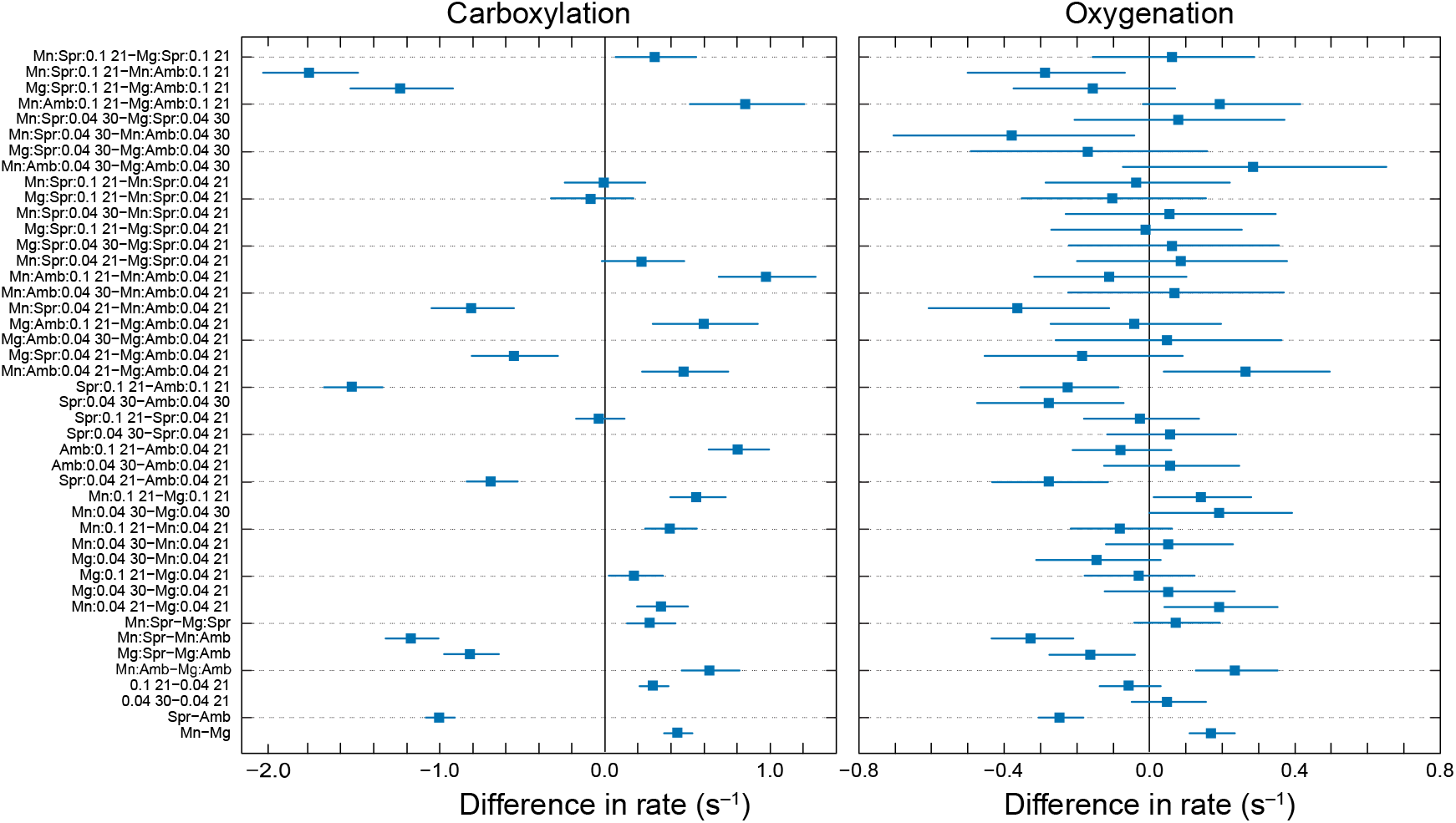
Tukey test comparing RuBP carboxylation and oxygenation rates under various combinations of metals (Mg vs. Mn), *Plantago* l*anceolata* ecotypes (Spr vs. Amb), CO_2_ concentrations (0.04% vs. 0.1%), and O_2_ concentrations (21% vs. 30%). Shown are mean difference ± SE, n = 4 –10.

The kinetics of the rubiscos from the two ecotypes differed more when they were associated with Mn^2+^ than Mg^2+^(**Fig. 3 & Table 1**). The rubisco from the ambient ecotype exhibited a slightly higher *V*_cmax_ and *V*_omax_ than the rubisco from the spring ecotype when bound to magnesium and a much higher *V*_cmax_ and *V*_omax_ when bound to Mn^2+^ (**Fig. 3**). Specificity for CO_2_ over O_2_ (*S*_c/o_) for rubiscos from both ecotypes were higher when the enzymes were bound to Mg^2+^ than Mn^2+^ (**Fig. 3**).

**Fig. 3.**
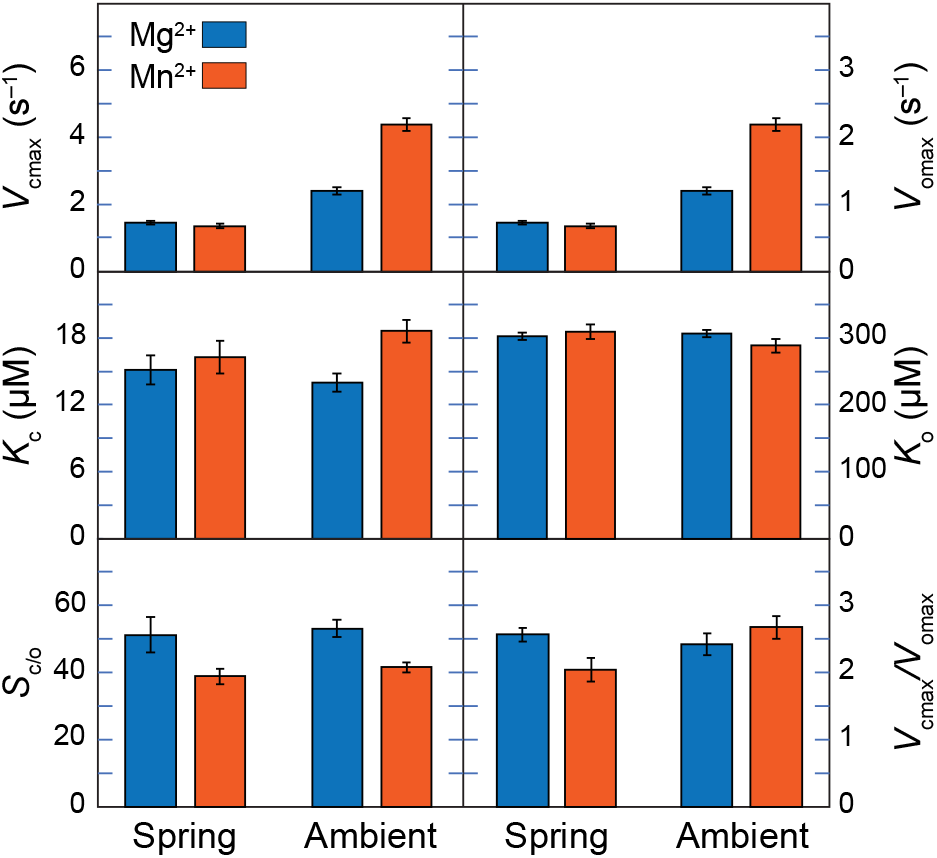
Influence of Mg^2+^ or Mn^2+^ (mean ± credible interval, n = 4 to 20) on the kinetics of rubisco purified from two ecotypes of *Plantago lanceolata* L.: one (Spring) collected near a CO_2_ spring has experienced high CO_2_ for many generations; the other collected nearby (Ambient) has experienced ambient CO_2_. *V*_cmax_ is the maximum velocity of carboxylation, *V*_omax_ is the maximum velocity of oxygenation, *K*_c_ is the Michaelis constant of rubisco for CO_2_, *K*_o_ is the Michaelis constant of rubisco for O_2_, *S*_*c/o*_ is the specificity of rubisco for CO_2_ over O_2_, and *V*_cmax_/*V*_omax_ is the ratio of the maximum velocities. Table 1 provides statistical analysis of these data.

**Table 1.** 2-way ANOVA for the influence of Ecotype, Metal, or their interaction on the kinetic parameters shown in Fig. 3 for rubiscos purified from the two ecotypes in the presence of Mg^2+^ or Mn^2+^. “Signif.” denotes significance where “.” *P* < 0.1, “*” *P* < 0.05, “**” *P* < 0.01, and “***” *P* < 0.001.

| Parameter | Factor | Deg. of freedom | Sum of squares | Mean of squares | F-value | P (>F) | Signif. |
| --- | --- | --- | --- | --- | --- | --- | --- |
| $V_{cmax}$ | Ecotype | 2 | 134 | 67 | 1089 | $<2 \times 10^{-16}$ | *** |
| | Metal | 1 | 4.43 | 4.43 | 71.9 | $2.6 \times 10^{-7}$ | *** |
| | Ecotype × Metal | 1 | 5.51 | 5.51 | 89.4 | $6.0 \times 10^{-8}$ | *** |
|  | Residuals | 16 | 0.99 | 0.06 |  |  |  |
| $V_{omax}$ | Ecotype | 2 | 20.8 | 10.4 | 2214 | $<2 \times 10^{-16}$ | *** |
| | Metal | 1 | 0.70 | 0.70 | 147 | $1.7 \times 10^{-9}$ | *** |
| | Ecotype × Metal | 1 | 0.38 | 0.38 | 80.0 | $1.3 \times 10^{-7}$ | *** |
|  | Residuals | 16 | 0.075 | 0.005 |  |  |  |
| $K_c$ | Ecotype | 2 | 4970 | 2485 | 407 | $1.9 \times 10^{-14}$ | *** |
|  | Metal | 1 | 40.1 | 40.1 | 6.57 | 0.021 | * |
| | Ecotype × Metal | 1 | 11.0 | 11.4 | 1.87 | 0.19 | |
|  | Residuals | 16 | 98.0 | 6.1 |  |  |  |
| $K_o$ | Ecotype | 2 | $1.8 \times 10^6$ | $9.2 \times 10^5$ | 3086 | $<2 \times 10^{-16}$ | *** |
|  | Metal | 1 | 249 | 249 | 0.84 | 0.37 |  |
| | Ecotype × Metal | 1 | 726 | 726 | 2.4 | 0.14 | |
|  | Residuals | 16 | 4747 | 297 |  |  |  |
| $S_{c/o}$ | Ecotype | 2 | $4.5 \times 10^4$ | $2.2 \times 10^4$ | 594 | $9.7 \times 10^{-16}$ | *** |
| | Metal | 1 | 780 | 780 | 20.6 | $3.3 \times 10^{-4}$ | *** |
| | Ecotype × Metal | 1 | 1.0 | 1.0 | 0.038 | 0.85 | |
|  | Residuals | 16 | 605 | 38 |  |  |  |
| $V_{cmax}/V_{omax}$ | Ecotype | 2 | 121 | 60.7 | 494 | $4.2 \times 10^{-15}$ | *** |
|  | Metal | 1 | 0.07 | 0.07 | 0.60 | 0.45 |  |
| | Ecotype × Metal | 1 | 0.82 | 0.82 | 6.67 | 0.020 | * |
|  | Residuals | 16 | 1.97 | 0.12 |  |  |  |

### cpDNA sequences

Both ecotypes had similar plastid DNA coding for the large unit of rubisco (rbcL). The rbcL gene is located at positions 53249 to 54682 in the ambient ecotype and 53018 to 54451 in the elevated ecotype of the chloroplast DNA sequences (cpDNAs). Gene maps showing the rbcL encoding regions and the translation directions are displayed in **Fig. 4**. The rbcL gene in the reconstructed sequences was identical to those in previously published sequences (Zhao et al. 2022, Saban et al. 2020). A small percentage of the fragments, however, differed from this sequence in one or, in one case, two nucleotides (**Table 2**). The most common of these, by a significant margin, was a change in nucleotide 890 from A to C, which occurred in approximately 5.6% of fragments in the spring ecotype and 4.1% in the ambient. This change replaces a methionine at residue 297 with a leucine.

**Fig. 4.**
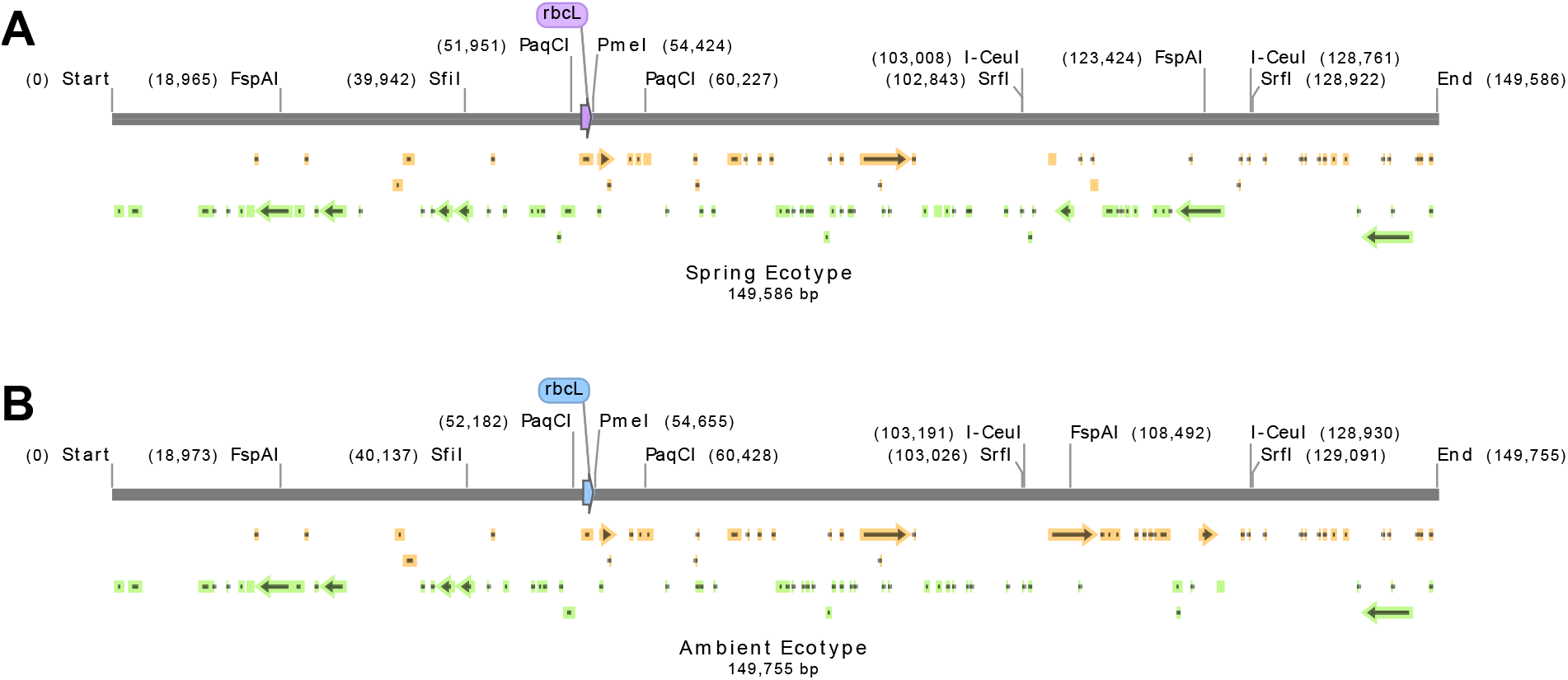
cpDNA map of rbcL from the (A) CO_2_ spring and (B) ambient ecotypes. The orange and green arrows indicate open reading frames (ORFs). The restriction enzymes shown are unique and dual cutters.

**Table 2.**
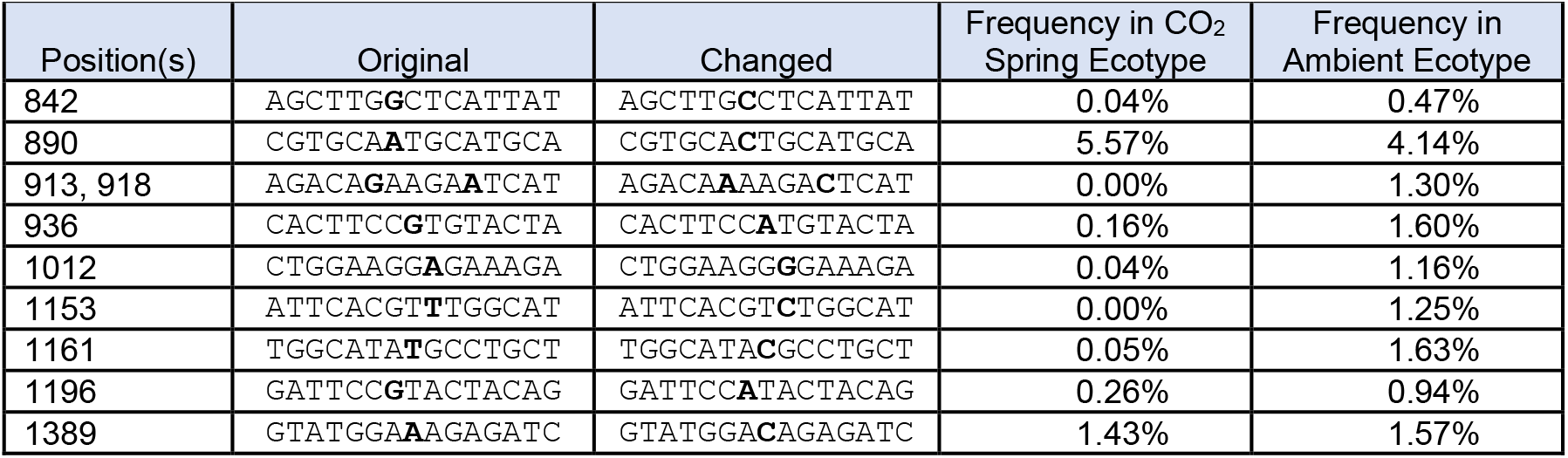
Mutations that appeared in a significant portion of the fragments matching the rbcL gene in two ecotypes of *Plantago lanceolata*. “Positions” refer to the positions of the changed nucleotide(s) and are specified relative to the start of the rbcL gene. “Frequency” refers to the number of fragments containing the changed sequence as a percentage of the total number of fragments containing the original sequence and the changed sequence.

## Discussion

### Kinetics

The ambient ecotype had overall higher rates of both carboxylation and oxygenation than the CO_2_ spring ecotype. One possible explanation is differences in genomic DNA. We noticed larger average seed size and lower seed yield per flower and per plant for the CO_2_ spring ecotype than for the ambient ecotype. *Plantago lanceolata* is able to change growth habit and phenology in different growth conditions; however, it is known that genetic variations exist based on significantly reduced seed yields compared to original and alien sites (Van Tienderen et al. 1991). Under limited water supply and high light exposure, *Plantago lanceolata* displayed higher water use efficiency (WUE), better nitrogen absorption, and higher bacterial frequency in roots (Miszalski et al. 2023). The photochemical activity of PSII stayed at the same level under the stressed conditions as under the no-stress conditions (Miszalski et al. 2023). Taken together, we think the extensive plasticity of *Plantago lanceolata* might be shared among many species from various geographic locations, and the variations in photosynthetic rates are genotype-specific.

Exposure to elevated CO_2_ atmospheres stimulated carboxylation rates in the ambient ecotype more than in the CO_2_ spring ecotype. This would not be expected if faster carboxylation rates were always beneficial. Were balancing carboxylation and oxygenation advantageous, then elevated and, especially, fluctuating levels of CO_2_ might select for a mechanism where carboxylation became less sensitive to atmospheric CO_2_. The nature of such a mechanism is unknown. The CO_2_ spring ecotype has adapted to elevated CO_2_ levels for many generations and might be expected to balance carboxylation and oxygenation more quickly. It is worth noting that the 0.1% CO_2_ treatment used in this study is generally not sufficient to saturate rubisco carboxylation, so the lack of significant change in the CO_2_ spring ecotype between ambient and 0.1% CO_2_ conditions may indicate an additional regulatory mechanism beyond the usual kinetics. The change from methionine at residue 297 with a leucine is too rare to have a significant effect on the overall kinetics. Possibly, the small subunit of rubisco, whose gene is located in the nucleus and was not sequenced in this study, has some regulatory effect.

In other C_3_ species that have been studied, carboxylation rates are higher when rubisco is bound to Mg^2+^, whereas oxygenation rates are higher when bound to Mn^2+^ (Shi et al. 2024). By contrast, manganese Mn^2+^ appears to promote both carboxylation and oxygenation in rubisco from *Plantago lanceolata*. Previous studies (Bloom and Kameritsch 2017) indicate that plants respond to higher levels of CO_2_ by increasing manganese levels. If rubisco from *Plantago lanceolata* thrives in the presence of Mn^2+^, this may explain why this species was able to adapt to the high CO_2_ environment of the spring.

When bound to Mg^2+^, kinetic parameters for rubisco from both ecotypes were roughly within the range of those for other C_3_ plants (Shi et al. 2024). Nonetheless, if the CO_2_ spring ecotype has a mechanism that preserves the balance of carboxylation and oxygenation under elevated CO_2,_ as introduced above, then Michaelis-Menten kinetics might not fully characterize it because Michaelis-Menten kinetics predict that the ratio of carboxylation to oxygenation should be proportional to the ratio of the two gases with *S*_*c/o*_ as the proportionality constant.

### Chloroplast DNA sequences

The reconstructed sequences for *rbcL* from *P. lanceolata* were nearly identical to the previously published sequences for the *rbcL* gene (Zhao et al. 2022, Saban et al. 2020). This supports the generality that the *rbcL* gene is highly conserved among species (Saban et al. 2020) and the *P. lanceolata-*specific findings of low overall genetic differentiation between the CO_2_ elevated and ambient ecotypes and weak selection in exons (Saban et al. 2020). Although DNA methylation may play a role in *P. lanceolata*’s evolutionary divergence under exposure to elevated and ambient CO_2_ for multiple generations (Saban et al. 2020), we did not conduct cpDNA bisulfite sequencing because chloroplast DNA is relatively insensitive to methylation (Fojitova et al. 2001, Ahlert et al. 2009) and because in particular *rbcL* genes in tobacco, *Arabidopsis*, peas, and tomatoes are not affected by DNA methylation (Fojitova et al. 2001). Moreover, the BstNI/EcoRII assay, which had been used in previous studies, might not be accurate for analyzing low levels of methylation in chloroplast DNA sequences. (Marano et al. 1991, Ngernprasirtsiri et al. 1988, Ohta et al. 1991, Langdale et al. 1991). Increasing adenine or cytosine methylation by introducing cyanobacterial genes for adenine and cytosine DNA methyltransferases into the tobacco plastid genome through chloroplast transformation resulted in no phenotypic differences from wild-type plants and no alterations in plastid gene expression (Ahlert et al. 2009).

The variant of the *rbcL* gene with a C, instead of an A, at position 890 occurred in both *P. lanceolata* ecotypes at a much higher frequency than other mutations (**Table 2**). This DNA change corresponds to a change from Met 297 to Leu 297 in the amino acid sequence, which we might understand by examining the related – and well-studied – spinach rubisco sequence. Rubisco crystal structures from spinach show residue 297 in close proximity to the substrate access channel (**Fig. 5**). In an activated transition state analog that was achieved by binding to the tight-binding inhibitor CAP, magnesium coordination is in a distorted octahedron with longer than usual metal-R bond distances (**Fig.5A**, Andersson 1996). This is the “closed” active site conformation. In another activated structure with the substrate RuBP and Ca^2+^, residue 297 moves to hover over the active site channel, whereas Lys334 moves away from the reaction center (**Fig. 5B**). Note that Ca^2+^ in **Fig. 5B** can activate rubisco but does not turn it over (Taylor and Andersson 1997). The active site appears to be “open”.

**Fig. 5.**
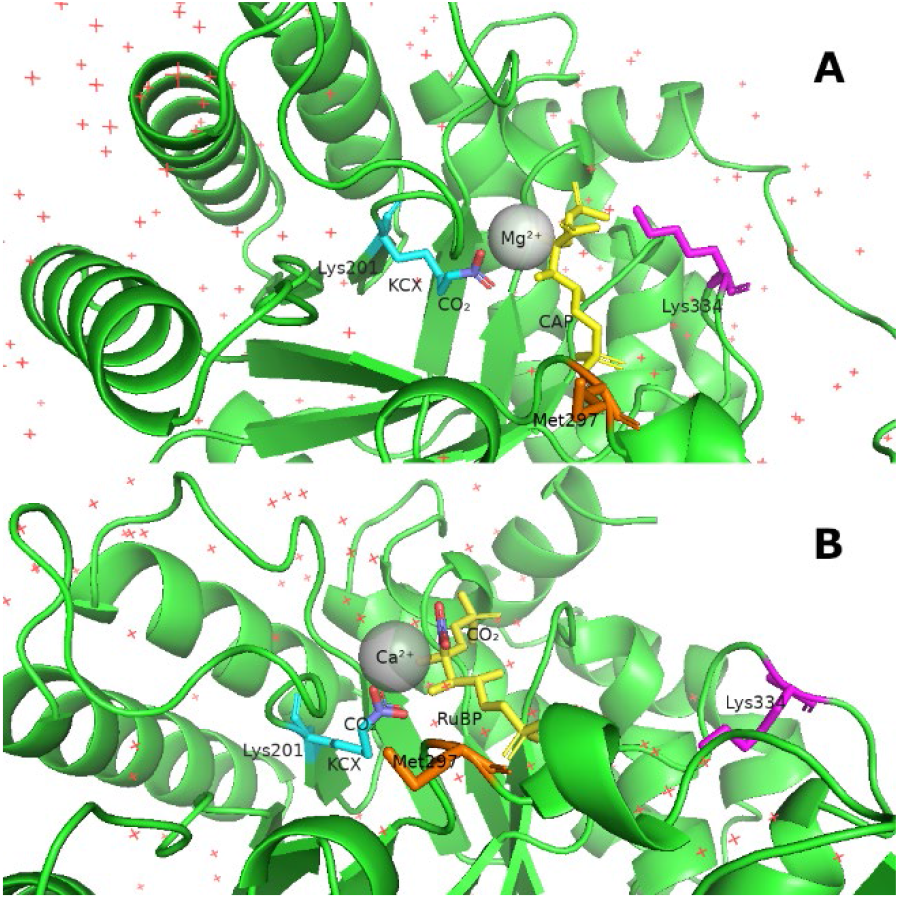
The vicinities of the active sites in two published crystal structures of spinach rubisco. The red “+” symbols indicate water molecules. (A) Rubisco bound to Mg^2+^ and CAP (PDB: 8RUC, Andersson 1996). (B) Rubisco bound to Ca^2+^ and RuBP (PDB: 1RXO, Taylor and Andersson 1997). KCX denotes lysine peptide linkage and CAP denotes 2-carboxyarabinitol-1,5-diphosphate.

Although interconversions of hydrophobic residues such as from methionine to leucine are common and enzyme function before and after such changes often remain the same, the significant location of residue 297 should not be ignored. It is located right at the entrance of the active site and at the end of the flexible part of a loop. Loop movements involving residue 297 might also interact with neighboring loops such as the loop containing the carbamylated Lys201 that is required for CO_2_ activation or the loop on the distal side of the substrate containing Lys334, whose interactions with substrate-CO_2_ complex are required for turning rubisco over (Cleland et al. 1998). Lys334 is also believed to stabilize the transition state and the product (Cleland et al. 1998) during the reactions when the active site is in the “closed” conformation (**Fig. 5A**). The distances of the C-alpha in Lys 334 to Met 297 changed from 20.6Å to 15.8Å from the open (**Fig. 5B**) to the closed state (**Fig. 5A**). The M297L variant encoded by this version of the gene might have properties that are advantageous under CO_2_ enrichment such as regulating the opening and closing of the active site access tunnel, which might result in chloroplasts with this variant being favored in the CO_2_ spring ecotype. Perhaps it occurred in the past and persists in a small percentage of chloroplasts in all ecotypes, but a high CO_2_ environment selects for an increase in that percentage.

As the most abundant protein on Earth, type I rubisco large subunit gene (*rbcL*) evolves slower than 98% of all enzymes (Bouvier et al. 2024). A phylogenetic study found that one nucleotide substitution and one amino acid change happen every 0.9 million years and 7.2 million years, respectively (Bouvier et al. 2024). This is consistent with the 4 to 5% occurrences of the mutation we observed at position 890 on the *rbcL* gene. Molecular dynamics simulations on extant and ancestral rubisco sequences showed that the small subunit (*rbcS*) consistently confined the structural dynamics of the large subunit throughout the enzyme’s evolutionary history (Kaustubh et al. 2024). The data were inconclusive, however, on the small subunit serving as a CO_2_ substrate reservoir (Kaustubh et al. 2024), as some have proposed (Van Lun M et al. 2014). This may be because each species contains only one copy of the *rbcL* gene in the chloroplast, whereas multiple *rbcS* sequences are found encoded in genomic DNAs, and *rbcS* evolves faster (Bouvier et al. 2024). This might also explain the kinetic differences between spring and ambient genotypes, but clearly, additional studies on the function of rubisco small subunit are required.

## Conclusions

The kinetic properties of rubisco from *P. lanceolata* generally align with those of other C_3_ plant species. Nonetheless, notable differences exist. In particular, association of rubisco with Mn^2+^, promotes faster carboxylation and oxygenation in *P. lanceolata* than association with Mg^2+^. This disparity may account for the species’ ability to acclimate to high CO_2_ environments. The *rbcL* gene, which is identical in both ecotypes except for a higher frequency of a rare variant in the spring ecotype, cannot explain differences between the two ecotypes.

Data on rubisco kinetics from non-model plant species remain limited, underscoring the need to examine the factors contributing to the differences between *P. lanceolata* ecotypes and among *P. lanceolata* and model species. Elucidating the role of metals Mn^2+^ vs. Mg^2+^ within the photosynthetic and photorespiratory pathways should enhance our understanding of plant adaptations to climate change.

## Author contributions

A.J.B. planned the research, obtained the funding, wrote the manuscript, and drew the figures and tables; X.S. developed the biochemical methods, edited the manuscript, and conducted experiments; N.M.H. also conducted experiments, performed the statistical analyses, and edited the manuscript.

## Acknowledgments

NSF grant CHE-19-04535 funded a portion of this research.

## Data availability statement

Data used in this work have been uploaded to Dryad and will be made public upon publication.

